# A Novel Hollow Fiber Infection Model (HFIM) for Antiviral PK/PD studies of CMV infection

**DOI:** 10.64898/2026.03.13.710048

**Authors:** Lalitya M. Sudarsono, Suzanne A.M. Wenker, Xuanlin Liu, Jorn Brink, Dirk-Jan van den Berg, J.G.C. van Hasselt, Anne-Grete Märtson

## Abstract

The hollow fiber infection model (HFIM) is a translational *in vitro* model that links time-varying human pharmacokinetic profiles to the associated viral dynamic responses, from which pharmacokinetic/pharmacodynamic (PK/PD) targets can be derived. Establishing such targets is essential for antiviral dose selection and optimization. This is particularly important for cytomegalovirus (CMV) infection treatment, which primarily affects vulnerable patient populations. PK/PD targets for ganciclovir, the first-line drug for treatment, are not yet defined. The lack of an undefined PK/PD target makes dose optimization challenging and may result in suboptimal exposure, prolonged toxicity, and the emergence of resistance. For the first time, we have demonstrated the use of a low-cost hemodialyzer hollow fiber cartridge with application for CMV infection using ganciclovir. We have established a system that 1) supports CMV culture for PD analysis, 2) reproduces a clinically relevant ganciclovir PK profile, and 3) maintains consistent drug exposure in the infected cells, allowing reliable PK/PD analysis. Quantitative methods such as tissue culture infectious dose 50% (TCID_50_) and quantitative PCR were used to assess both active virus replication and genome copies production. Ganciclovir PK was measured using liquid chromatography-tandem mass spectrometry (LC-MS/MS). This validation study serves as a fundamental step that can allow further PK/PD studies for ganciclovir and other antiviral agents that is still largely understudied. Consequently, this model could provide an affordable and practical platform for establishing clinically relevant PK/PD targets and guide treatment optimization.

## Introduction

Pharmacokinetics/pharmacodynamics (PK/PD) plays a central role in anti-infective drug development and treatment optimization. PK/PD is essential to understand how, for a given compound, the exposure dynamics over time can be linked to its viral dynamic outcomes. In antibacterial therapy, the PK/PD for several antibiotics and antifungals such as glycopeptides, aminoglycosides and triazoles, is well defined. This understanding refers to whether a compound is concentration-dependent, time-dependent, and/or both concentration and time-dependent, to achieve an optimal therapeutic target. It is, however, still unclear whether the same principles apply for antivirals, as the exposure-response relationships (PK/PD) have not yet been well characterized nor established (1).

The HFIM is an *in vitro* system that has been used to reproduce human PK and evaluate the expected PD response. While commonly used for antibiotic studies, the use of HFIM is less common in antiviral studies (2). Mimicking the human PK is critical as drug clearance and exposure over time can have a significant influence on viral load decline (1, 3, 4). These variables cannot be mimicked in conventional *in vitro* assays, where static concentrations are maintained over time and clinical dosing regimens cannot be tested (5–7). Currently, there are limited PK/PD studies for antivirals available. This is particularly evident in the case of cytomegalovirus (CMV) infection treatment, where PK/PD targets have not been well established (8, 9).

CMV is a latent herpesvirus that causes severe diseases and high mortality rates in immunocompromised individuals, which are often reported in transplant recipients (10). Ganciclovir and its prodrug, valganciclovir, are the first-line drug of choice for CMV treatment in clinical practice (11, 12). Treatment with ganciclovir is challenging due to undefined therapeutic targets and high frequency of clinical toxicity (8, 13, 14). Alternative anti-CMV drugs like maribavir have been used to tackle toxicity, but PK/PD targets are yet to be defined (15). Given the limited PK/PD understanding for current anti-CMV therapies, there is an urgent need to define such targets to optimize CMV treatment and achieve better clinical outcomes (14–18).

In this study, we aimed to establish and characterize a robust CMV HFIM that can support CMV infection while also reproducing a human-like PK profile of ganciclovir. Our main objective was to provide a reproducible and affordable framework for PK/PD assessments of ganciclovir, which can be extended for other antiviral agents, with the use of a low-cost hemodialyzer.

## Results

### CMV replicates actively in the established HFIM

We established the hollow fiber-CMV infection set-up as schematically presented in **Figure 1**, further details on culturing conditions and the framework of the set-up are described in Methods. The hollow fiber cartridge is used to support CMV infection, where a continuous flow of the cell culture growth media from the central reservoir is maintained throughout the cartridge. It serves as an infection site, where the human lung fibroblasts (MRC-5) cells and CMV strains were inoculated and isolated. To evaluate whether our system can support CMV infection, we used two strains; the Towne, a lab adapted strain and the BAC-derived, Merlin strain, which is genetically similar to the wild-type strain (19). Consequently, we observed a 3-and 4-fold log_10_ increase of genome accumulation (copies/mL) for Towne and Merlin strain respectively. Meanwhile, a 4-and 10-fold log_10_ growth of active virus titers (PFU/mL) was observed for the Merlin and Towne strains, respectively, over 6 days of infection period (**Figure 2**.). The system could, therefore, support a long-term infection period, which can be used for further PD assessments.

**Figure 1.**
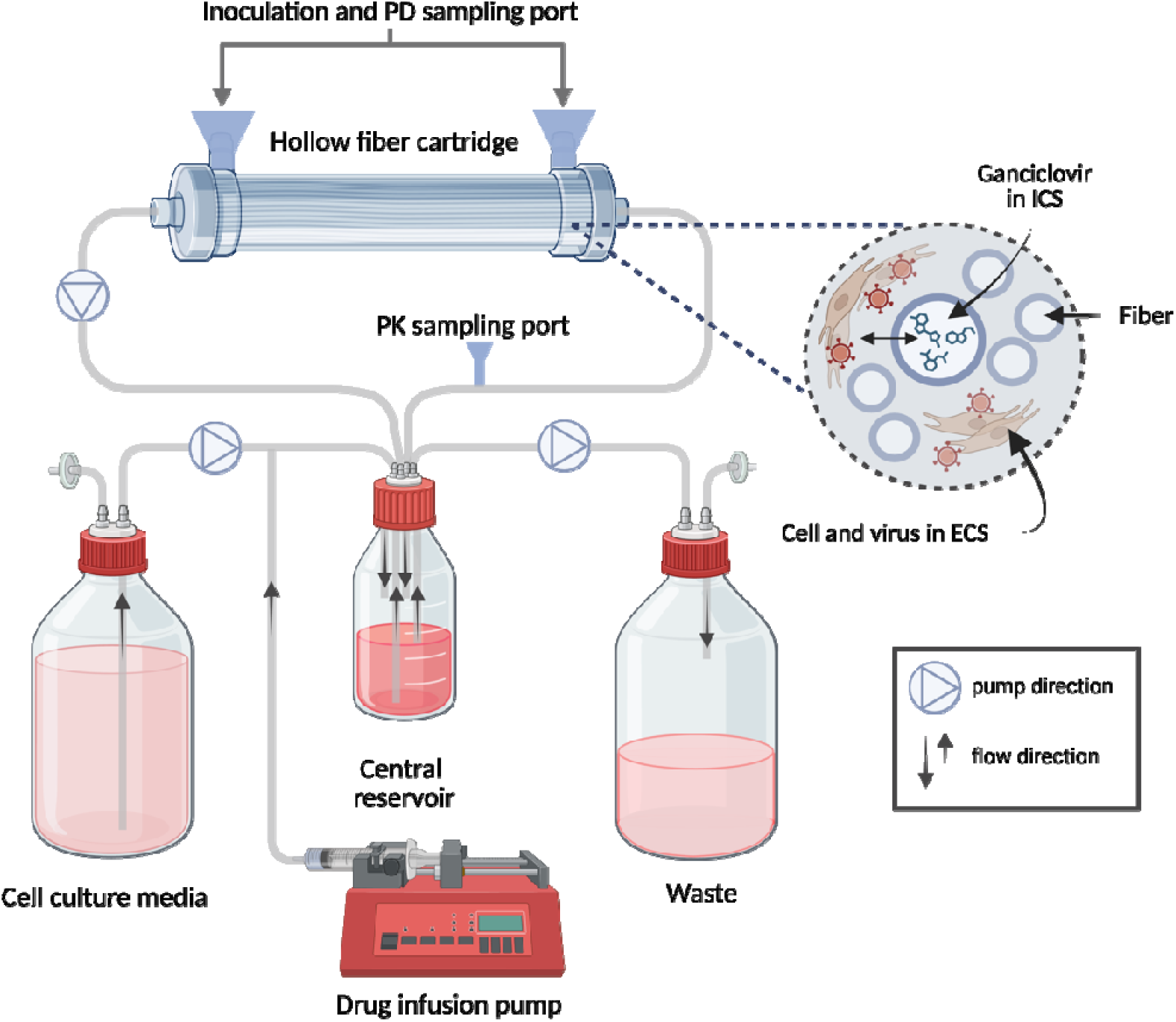
Schematic overview of the hollow fiber infection model. The hemodialyzer cartridge inoculates MRC-5 cells and CMV in the extra-capillary space (ECS). A continuous flow of cell culture media into the cartridge was maintained from the central reservoir. For drug treated system, drug was infused directly into the central reservoir and flowed along the intracapillary (ICS) of the hollow fiber. To mimic drug clearance, an inflow of drug-free cell culture media was set accordingly into the central reservoir, and the surplus was flowed out into the waste. Cartridge and central reservoir were maintained at 37° C and 5% CO_2_ during the experiment.

**Figure 2.**
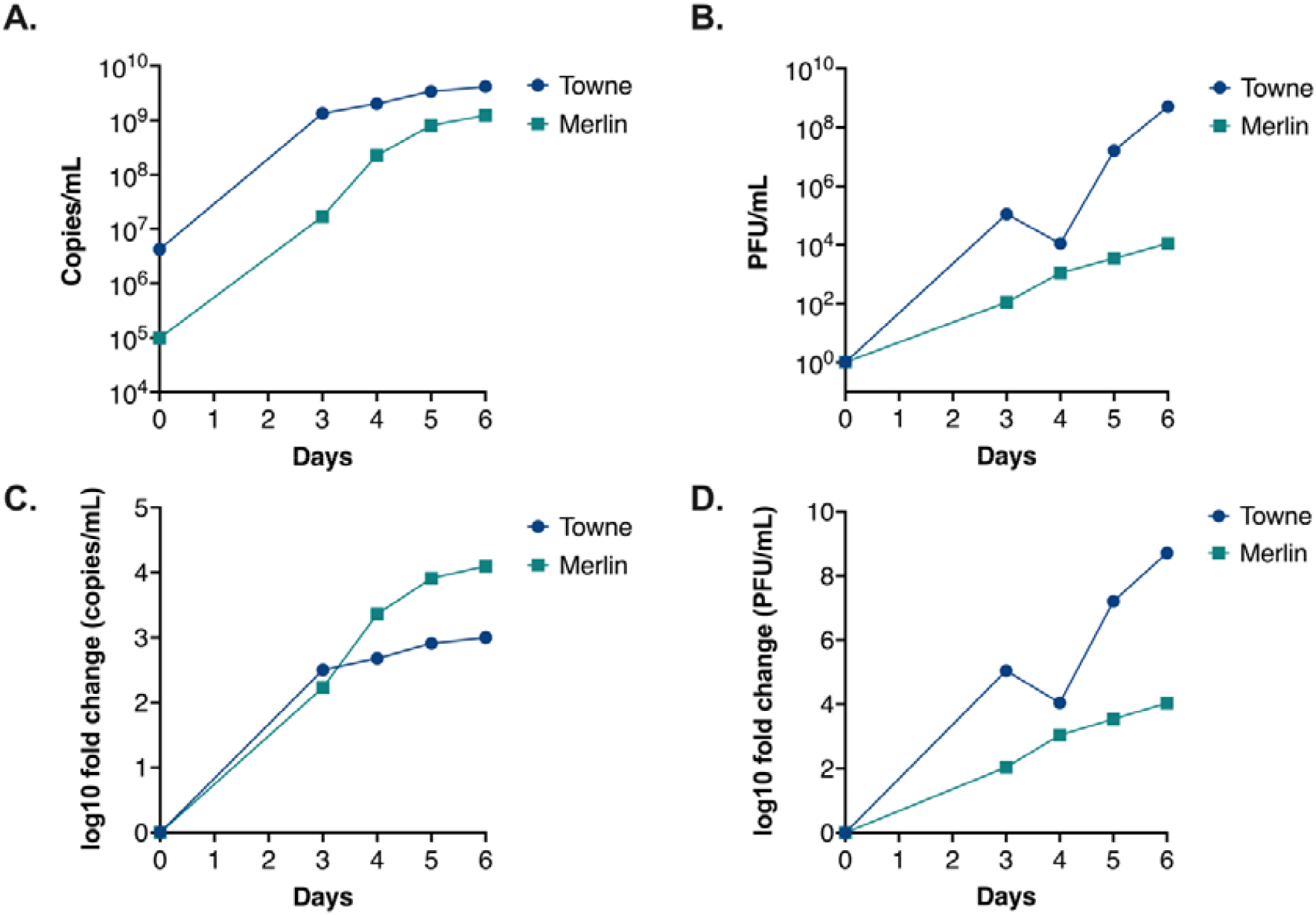
CMV replicates in MRC-5 cells within the HFIM system. Circle and square lines represent the Towne (lab-adapted) and the Merlin (wild-type) strain, respectively. (**A, B**) represent the genome copies (copies/mL) and virus titer (PFU/mL), over time. (**C, D**) represent the log_10_ fold change relative to quantified copies/mL and PFU/mL at day of inoculation.

### HFIM reproduces human-like ganciclovir PK plasma profile

Ganciclovir was infused directly into the central reservoir, as represented in **Figure 1**. at 2 mg/h (continuous infusion), administered every 12 h, as in clinical practice. To evaluate how well the HFIM can reproduce a human-like PK, we simulated expected PK curves based on the input parameters. The observed and the simulated concentration-time profile were compared, and similar profiles were achieved (**Figure 3**.). The observed data were fitted to a one-compartmental model. Thereby, the clearance, volume of distribution, elimination constant rate (K_e_), and 12h area under the concentration-time curve (AUC_0-12h_) were also estimated using a one-compartmental model fitting (**Table 1**.). We have observed an overall good fit of the observed PK profiles. Therefore, clinical dosing regimens of ganciclovir can be performed in the established HFIM.

**Table 1.** Input vs. estimated ganciclovir PK parameters of the HFIM with one-compartmental fitting analyzed in R.

| PK parameters | Input | Estimates (%RSE) |
| --- | --- | --- |
| Clearance (L/h) | 0.03 | 0.033 (0.2) |
| Volume of distribution (L) | 0.2 | 0.3 (2.05) |
| Elimination rate constant (h<sup>-1</sup>) | 0.15 | 0.11 |
| AUC<sub>0-12</sub> (mg.h/L) |  | 42.2 |

**Figure 3.**
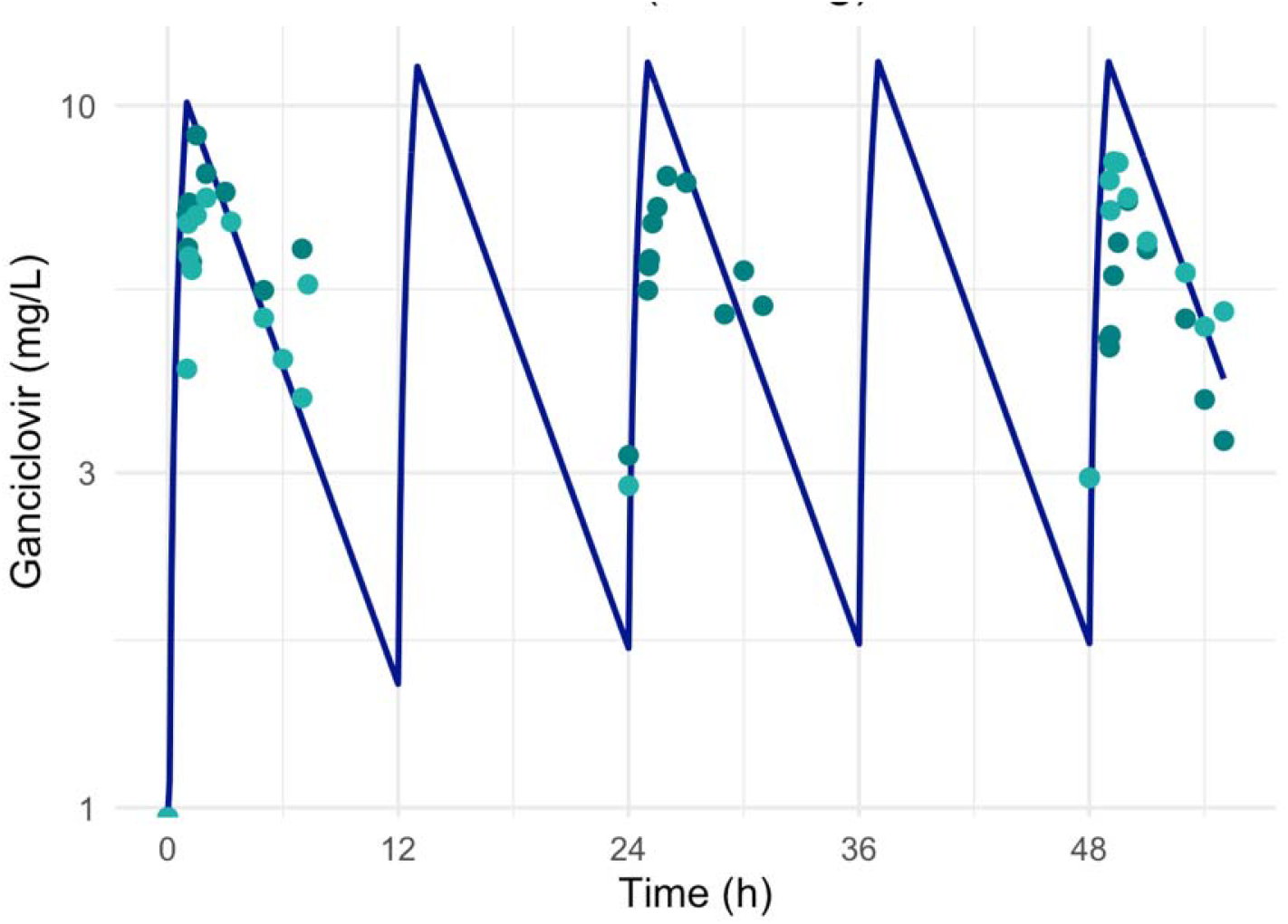
Concentration-time profile of observed and simulated (line) ganciclovir concentrations. Blue and teal dots represent data points from two independent experiment.

### Ganciclovir maintains active exposure into infection site without binding in the HFIM system

A high flow rate (30 mL/min) was set up to achieve a constant equilibrium between ganciclovir concentrations in the central reservoir, the extra-capillary space (ECS) and the intra-capillary space (ICS) of cartridge. To validate whether this is consistent over time, we sampled media containing drug from the ECS which isolates the virus and cells, and the ICS, which comes directly from the central reservoir (Figure 4A). We have observed that the ganciclovir PK curve of the cartridge is similar to that of the central reservoir (Figure 4B.) Throughout different sampling time points, we observed no significant difference between ganciclovir concentrations as demonstrated by the estimation plot (Figure 4C). No significant differences were observed for the calculated AUC_24h_ between the two sampling points (Figure 4D). These results demonstrate that the system supported an active diffusion between the ECS and ICS, while maintaining constant equilibrium between the concentrations in the central reservoir and those to which the cells and viruses were exposed. This suggested that the compound did not accumulate in the cartridge and no drug binding was present in the system, allowing direct exposure to the infected cells. Overall, indicating that the time-varying drug exposure within the infection site can be evaluated with the eventual PD response.

**Figure 4.**
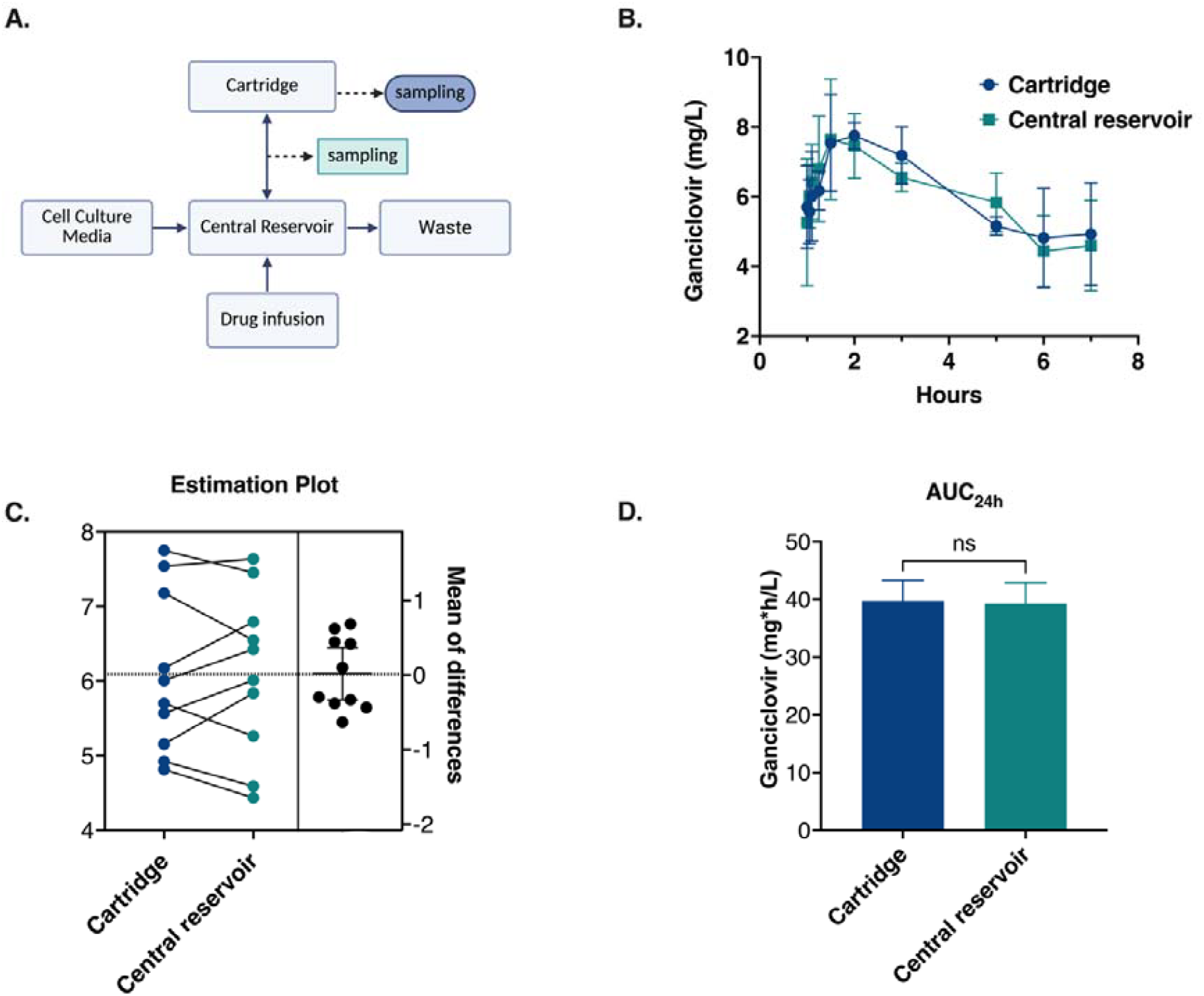
Ganciclovir concentrations in cartridge (ICS) vs. central reservoir (ECS). **(A)** Schematic of the HFIM system showing the two sampling ports. (**B**) Ganciclovir concentration profile. (**C**) Differences and mean of differences in concentrations between cartridge and central reservoir sampled at different time points. (**D**) AUC_24h_ of ganciclovir concentration in the cartridge and central reservoir of the HFIM (P-value>0.05, N=3).

### Pharmacodynamics of CMV during ganciclovir exposure

To characterize PD response under ganciclovir exposure, we administered ganciclovir after three days of viral growth. We then evaluated viral production of both Towne and Merlin strains in terms of the DNA and virus titer replication for 72 hours post-treatment. For this, we assessed the log_10_ fold difference relative to the copies or titer observed at day 0, when treatment was initiated. For Towne, we have observed full absence of growth for both total copies/mL and virus titer throughout 72 hours of exposure compared to the control arm **(Figure 5A, 5C)**. For Merlin, we initially observed an increase for both copies and virus titer after 24-hour post-treatment, thereafter, absence of growth for were observed **(Figure 5B, 5D)**. Under exposure of ganciclovir at 2 mg/hour, q12-hour, we have observed an overall absence of growth in genome copies and virus titer throughout 3 days post-treatment initiation.

**Figure 5.**
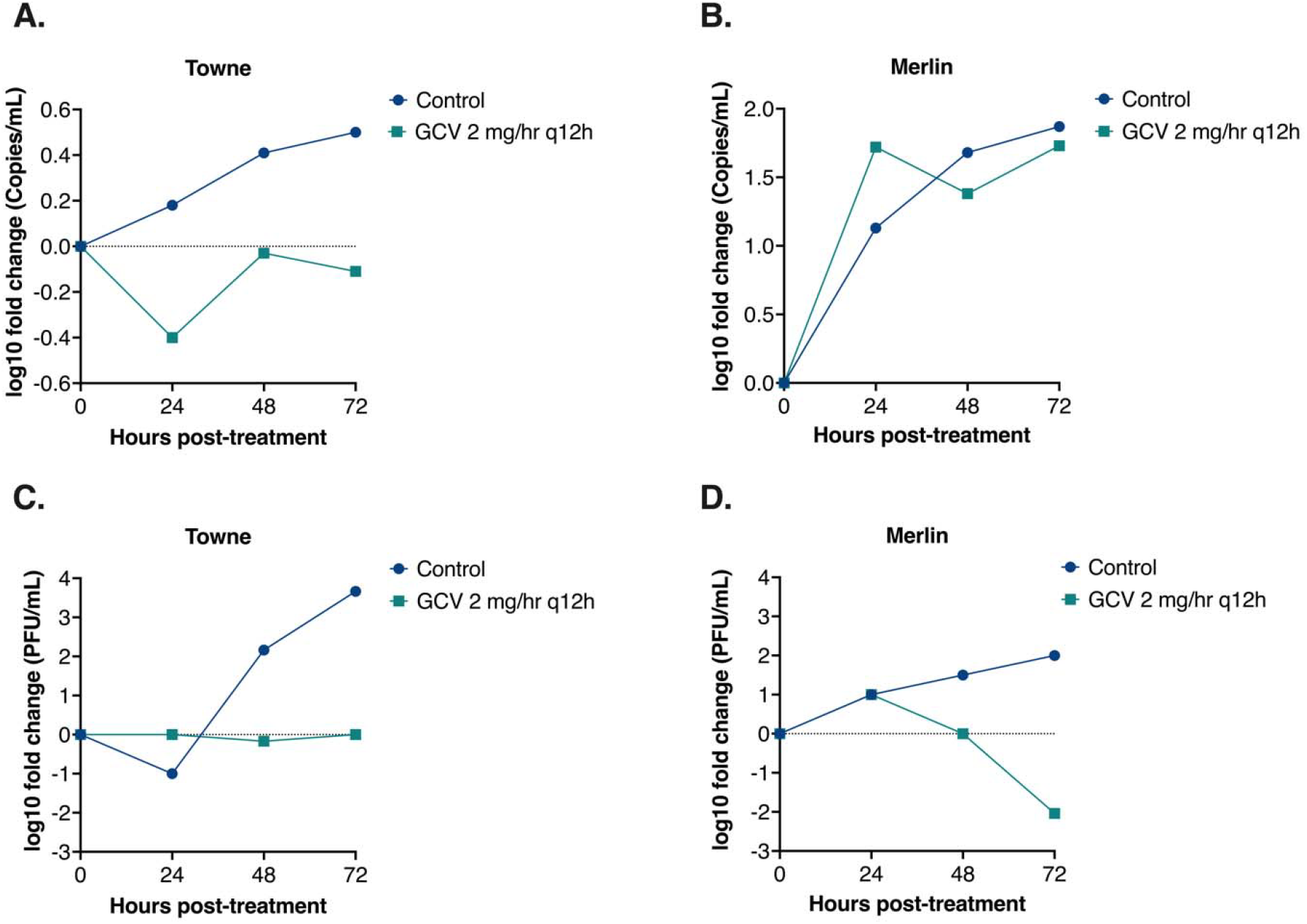
CMV growth with and without ganciclovir treatment in HFIM. Ganciclovir treatment was administered 2 mg/hour every 12 hours, 3 days post-inoculation. Circle and square lines represent the control (no drug treatment) and drug treatment group. **(A-D)** represent the log10-fold change of genome copies (copies/mL) and virus titers (PFU/mL) for Towne and Merlin, respectively, over 72 hours post-treatment initiation.

## Discussion

Here, we present for the first time, an established low-cost hollow-fiber system that can be applied for ganciclovir treatment-optimization studies. Our set-up can support a long-term CMV *in vitro* culture of both lab-adapted and wild-type strains, which can allow pharmacodynamic evaluations. The system might also, therefore, be used to amplify different CMV strains in a large volume, which might be useful for infection studies. Importantly, we have reproduced a concentration-time profile for ganciclovir that mimics the human plasma concentration profile where the estimated AUC_0-12h_ is within the expected clinical target range (40-60 mg.h/L) (20, 21). The result as well confirmed direct ganciclovir exposure to CMV-targeted infected cells within the hollow-fiber cartridge. This means that there was no delay of drug diffusion and distribution between the intracapillary space (ICS) and the extracapillary space (ECS). Consequently, we have observed virus growth suppression when a clinical-based dosing regimen was applied over 3 days post ganciclovir treatment initiation. Altogether, these results have demonstrated the practicality of the setup for further PK/PD assessments of ganciclovir in CMV infection therapy.

Furthermore, an important advantage of our HFIM system is the use of a hemodialyzer as a low-cost hollow fiber cartridge (up to 25€). The commonly used commercial cartridges are known to be costly (up to 1000€), which can limit accessibility for academic research. The costs can increase exponentially for cell culture, wherein a single-use cartridge is often used for every experimental arm. Recent studies have shown the feasibility of the hemodialyzer as a cost-effective hollow fiber cartridge for several antibiotic studies (22–25), however, this has never been demonstrated for antivirals. For antiviral studies, the experimental condition is complex as both the host cell and virus culture need to be maintained inside the cartridge. Our set-up has shown the feasibility of viral culture and might therefore be applicable for PK/PD studies of other antiviral agents. However, the culturing conditions in the cartridge, such as the cell lines, the cell number, the virus strain, the multiplicity of infection (MOI), as well as the effect of drug binding within the system, might require further optimization.

The use of an infection model with a dynamic drug exposure set-up is critical for establishing clinically relevant therapeutic targets. Static drug concentration-based assays are limited by the lack of drug clearance and thus clinical dosing regimens are difficult to study (26–28). The established HFIM in this study provides a validated *in vitro* set-up where these limitations can be mitigated. The viral dynamic outcomes can then be evaluated depending on the drug exposure profile over time. This is critical for PK/PD studies of antiviral agents where exposure-response parameter relationships are still understudied (15, 29). Especially for CMV infection treatment, where the therapeutic window is difficult to define as special population groups are impacted (14, 18). This concerns mostly transplant recipients, who are immunocompromised and thus vulnerable to invasive sampling procedures in clinical studies. In the case of ganciclovir, current standard dosing is currently applied across various patient profiles, which has led to a sub-exposure, slow decline of viral load suppression and clinical toxicity (8, 16–18, 30). There is thus an urgent need to optimize therapeutic targets and evaluate the exposure-effect relationship of ganciclovir in CMV treatment.

The established HFIM can be used essentially to perform dose-ranging and dose-fractionation studies. Previous studies have used the system to optimize and define therapeutic targets in cases of Influenza A, Zika and HIV infection treatments (3, 31–33). This is especially useful in the light of the recurring emergence of viral resistance (34, 35). Additionally, the infection model also serves as an advantage when recreating infections in immunocompromised conditions. While the HFIM model can offer a human-like drug-exposure profile on a cellular level, current state-of-the-art organ-on-a-chip systems might offer an alternative understanding of PK/PD evaluations on a tissue level and across multiorgan systems (36).

This study has some limitations. The HFIM allows controlled drug concentration exposure, which is beneficial predominantly for studying exposure-response relationships. However, the interplay between infection and immunity response on the PK/PD outcomes needs to be further evaluated and validated using additional experimental set-ups. The role of CMV-specific CD4^+^ and CD8^+^ T cells in controlling CMV infection needs to be considered when translating results into the clinical setting (37, 38). Incorporating these host immune parameters into a viral dynamic model may however, mitigate these limitations (39).

Finally, this infection model will ultimately support data generated from animal models and clinical trials, thereby paving ways to accelerate PK/PD studies and dose finding. Fundamental understanding of the relationship between ganciclovir exposure and viral dynamics is an essential step for defining clinically relevant targets. Such targets can be evaluated systematically using the HFIM system, which can be applied across multiple stages of drug development in addition to being cost-effective.

In conclusion, we developed a novel and affordable dynamic *in vitro* model for antiviral treatment optimization studies. The results of our study have shown the feasibility for further PK/PD assessments of ganciclovir in the established HFIM set-up. This is especially useful to define and optimize clinical targets of ganciclovir in CMV infection treatment.

## Material and Methods

### Cells and virus

MRC-5, human lung fibroblast cells (ATCC^®^ CCL-171™) were cultured and maintained at 37° C with 5% CO_2_, in Minimum Essential Medium (MEM; Capricorn Scientific) supplemented with 10% fetal bovine serum (FBS; Sigma-Aldrich^®^), 1% GlutaMAX™ - I CTS™ (Gibco^®^), 1% non-essential amino acids (NEAA; Gibco^®^), and 1% penicillin-streptomycin (Gibco^®^). Trypsin (Capricorn Scientific) was used for passaging and collection of adherent cells. Human cytomegalovirus (HCMV) Towne (ATCC^®^ VR-977 ™) and the Merlin strains were used for the infection model. The Merlin strain was originated from BAC-derived clones which are genetically similar to the Merlin (wild-type) strain (19). This was kindly provided by Professor Richard Stanton, from the Virology Immunology group, Cardiff University. The two strains were propagated in MRC-5 cells and viral stocks are used for infection. Viral stock titers were quantified by performing the TCID_50_ assay. An MOI of 0.1 (Towne) and 0.01 (BAC) is used for infection experiments.

### Hollow-fiber infection model

#### a. Cell culture and virus infection

MRC-5 cells were maintained and expanded in filtered cell culture flasks until confluency is reached, and concentrations were reached for experiment. 5.0 x 10^7^ cells were collected, and virus were injected altogether into the cartridge through the inoculation ports with syringes. Media containing cells and viruses were mixed carefully from the two ports to allow an even distribution throughout the hollow fiber cartridge. Thereafter, inoculated cells and virus were incubated at 37° C with 5% CO_2_ for 3 days to allow settlement and growth of cells and virus.

#### b. General overview of the HFIM system

A diagram of HFIM system is presented in Figure 1. The main circulation loop connects the central reservoir and the dialysis cartridge (FX paed hemodialyzer, Fresenius Medical Care, Bad Homburg, Germany), via a peristaltic pump (DT15, BT100F-1, Lead Fluid). The hollow fiber is represented by the cartridge that contains semi-permeable polysulfone (Helixone^®^) membranes (fibers) with a pore size of 3.3 nm, separating the intracapillary (ICS) and the extra-capillary space (ECS). This allows diffusion of drug compounds between the ICS and the ECS while keeping the cells and viruses retained in the ECS. The drug is administered with a syringe pump (ALADDIN-1000, World Precision Instruments) that infuses towards the central reservoir. To mimic the drug half-life (clearance), a second peristaltic pump was used to add drug-free cell culture media to the central reservoir. The third peristaltic pump was set at a slightly higher rate, discarding the excess volume to the waste. The synergy of the two pumps dilutes the drug while maintaining a constant volume of media in the central reservoir. Silicone and polypropylene-based tubes were used to circulate the fluid. Tubes and connections were sterilized beforehand, and the final set-up was done under the flow hood to maintain sterility.

The flow rate in the main circulation loop was set to 30 mL/min to achieve a rapid equilibration of drug concentrations in the central reservoir and, between the central reservoir and the ICS of the cartridge. An automated drug infusion program was used for drug administration where different dosing regimens can be applied: https://github.com/LeidenPharmacology/LeidenSyringePump.

Before cells and virus were inoculated, the system was primed with DPBS (1×) with Mg^2+^ and Ca^2+^ (Capricorn Scientific) to allow better cell attachment. Afterwards, priming was followed with complete growth media, overnight. Then, media containing cells and viruses were introduced via the inoculation port. Sampling of viruses were done with syringes that connects to the two ports that are connected to the ECS of the HF cartridge. Media containing drug samples were collected with syringes through a three-way port between the HF cartridge and the central reservoir.

#### c. PK parameters estimation and hollow fiber set-up reproduce human plasma PK profile

To reproduce human plasma PK profile, we targeted 10 mg/L as the peak concentration (C_max_) value, which corresponds to the range observed in patients (40). With the current dosing recommendation at 1-hour infusion (t), the drug infusion rate (K_0_) was calculated based on the following equation (multiple dose):

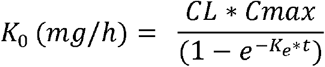

For all targeted C_max_ values, the elimination constant was calculated based on a typical half-life (t_1/2_), which was assumed to be 4 hours, where:

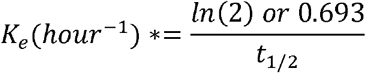

Clearance was calculated based on K_e_ and the volume of central reservoir, which was kept constant at 0.2 L,

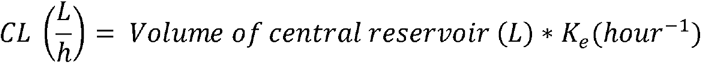

Ganciclovir (Thermo Fischer Scientific Chemicals) was freshly prepared at 1 mg/mL in DMSO and working solution was prepared and sterile filtered at 250 mg/L in water. Drug infusion rate (R) was set according to the dose (mg/h) where administered every 12 hours. Drug infusion pump rate at R was calculated according to:

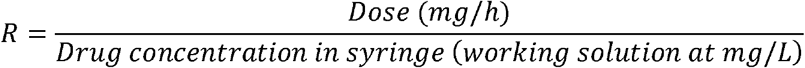

### Pharmacokinetic analysis of ganciclovir

Samples collected were stored -80° C until ready to be measured by the UHPLC-MS/MS. GCV-D5 was used as an internal standard (Toronto Research Chemicals, Toronto, Canada, G235002). The Shimadzu Nexera X3 UHPLC (Kyoto, Japan) coupled to a SCIEX 6500+ QTRAP mass spectrometer with an electrospray ionization (ESI) source (Sciex, Malborough, MA, USA) was used for our bioanalysis. A Kinetex LC Column (Phenomenex, 1.7 µm HILIC 100 Å, 50 × 2.1 mm) integrated with a security guard holder for the column (Phenomenex, 2.1 to 4.6mm ID) were employed for chromatographic separation. The temperature of column oven was kept at 30 ℃. Two sets of mobile phase combinations were used for bioanalysis. 0.1% formic acid and 1 mM ammonium formate served as aqueous phase (mobile phase A), meanwhile, 99.75% of acetonitrile, 0.1% formic acid and 1 mM ammonium formate was employed as an organic phase (mobile phase B) to elute ganciclovir. Acetonitrile, methanol and formic acid were of LC-MS quality (Biosolve BV, Valkenswaard, the Netherlands). Prior to measurement, all samples were spiked with internal standard and are diluted in methanol, thereafter, mixed and centrifuged for 10 minutes, 20.000× g at 4° C. Final dilution of supernatant at 1/1000 were prepared for measurement. A standard curve of ganciclovir was used quantify samples where 1/x^2^ is equal to or more than 0.99. Three quality control points (0.2, 4, and 40 µg/mL) were used to validate the standard curve. The gradient LC program started from 100% B for 0.5 min, increased to 50% B and held for 2.5 min, returned to 100% B, and equilibrated for 2 min. The run time was 5.5 min with the 0.3 mL/min of flow rate. SCIEX OS software platform (version 3.0) was used for acquisition and qualification processing.

### PK analysis

The observed concentrations sampled from the ICS were computed for fitting estimations. The expected PK profile was simulated using rxode2 package. The estimation of PK parameters was fitted using a one-compartmental model with the nlmixr/nlmixr2 package in R software (2024.12.1+563).

### DNA extraction and quantitative PCR for viral quantification

Extracellular viral samples were treated with a recombinant proteinase K (Thermo Fischer Scientific™) at 65⍰°C for 30 minutes, followed by heat inactivation at 90⍰0°C for 10 minutes. DNA was precipitated with isopropanol and washed with 75% ethanol. DNA pellets were airdried and resuspended in 30 µL of nuclease-free water. DNA extracts were stored at -20 °C. qPCR was performed using the primers targeting the UL83 gene in 96-well fast plates (Applied Biosystems™) with the following mix per sample: SensiMix SYBR low-ROX (QT625-05, Meridian BioScience), forward primer ((5’-3’): GGCTTTTACCTCACACGAGCATT) and reverse primer ((3’-5’): GCAGCCACGGGATCGTACT), NH_4_Cl buffer (10×, GC Biotech), and MgCl_2_. Plates were and run using standard 40-cycle, ABI 7500 system. Standard curves were generated from a 10-fold serial dilution of quantification CMV genomic DNA (VR-538 ™, ATCC).

### Viral quantification with TCID_50_

MRC-5 cells were seeded at a cell density of 1.5×10^4^ cells/well in 96-well plates for 24 h, at 37°C with 5% CO_2_. Tenfold serial dilutions of the viral samples were performed in 6 replicates (20 μL/well). Cytopathic effect (CPE) of each well was evaluated where positivity is defined when less than 25% of adherent cells were observed. TCID_50_/mL and the predicted value of PFU/mL (conversion factor 0.7) were calculated with TCID_50_ calculator developed by Marco Binder, University of Heidelberg, based on Spearman - Kärber method.

### Statistical analysis

Visual plots and statistical analysis were performed using the PRISM10 GraphPad and R Software. To visualize statistical differences, the mean ± SD is reported. To assess statistical significance between drug concentrations samples from the ICS and ECS, we firstly performed normality tests and then a paired t-test. A P-value of ≤0.05 was significant.

## Acknowledgements

We would like to thank Professor Richard Stanton that has kindly gifted us the Merlin BAC-derived strains. Figure 1 and 4A. were created in https://BioRender.com. AGM is funded by the Dutch Research Council (NWO), project ‘A Fresh Look at Antiviral Treatment: A Translational Pharmacology Approach’ with file number 09150162310191 of the research programme VENI.

## Notes

### Competing Interest Statement

AGM is funded by the Dutch Research Council (NWO), A Fresh Look at Antiviral Treatment: A Translational Pharmacology Approach, with file number 09150162310191 of the research programme VENI.

